# Correlative transmission electron microscopy and high-resolution hard X-ray fluorescence microscopy of cell sections to measure trace elements concentrations at the organelle level

**DOI:** 10.1101/2020.11.21.392738

**Authors:** Vanessa Tardillo Suárez, Benoit Gallet, Mireille Chevallet, Pierre-Henri Jouneau, Rémi Tucoulou, Giulia Veronesi, Aurélien Deniaud

**Affiliations:** ESRF, The European Synchrotron. 71 avenue des Martyrs, 38000 Grenoble, France; Institut de Biologie Structurale, CEA, CNRS, Univ. Grenoble Alpes, 71 avenue des Martyrs, F-38042 Grenoble, France; Univ. Grenoble Alpes, CNRS, CEA, IRIG, Laboratoire de Chimie et Biologie des Métaux, F-38000 Grenoble, France; Univ. Grenoble Alpes, CEA, IRIG, MEM, F-38000 Grenoble, France

**Author notes:** These authors contributed equally to this work (Vanessa Tardillo Suárez, Benoit Gallet). Correspondence should be addressed to **Dr Giulia Veronesi** or to **Dr Aurélien Deniaud**.

**Keywords:** X-ray fluorescence microscopy, Transmission electron microscopy, Hepatocytes, Mitochondria, Silver.

## Abstract

Metals are essential to all forms of life and their concentration and distribution in the organisms are tightly regulated. Indeed, in their free form, metal ions are toxic. Therefore, an excess of physiologic metal ions or the uptake of non-physiologic metal ions can be highly detrimental for the organisms. It is thus fundamental to understand metals distribution and dynamics in physiologic or disrupted conditions, for instance in metal-related pathologies or upon environmental exposure to metals. Elemental imaging techniques can serve this purpose, by allowing the visualization and the quantification of metal species in a tissue or down to the interior of a cell. Among these techniques, synchrotron radiation-based X-ray fluorescence (SR-XRF) microscopy is the most sensitive to date, and great progresses were made to reach spatial resolutions as low as 20×20 nm^2^. Until recently, 2D XRF mapping was used on whole cells, thus summing up the signal from the whole thickness of the cell. In the last two years, we have developed a methodology to work on thin cell sections, in order to analyze the metal content at the level of the organelle. Herein, we propose a correlative method to couple SR-XRF to electron microscopy, with the aim to quantify the elemental content in an organelle of interest. As a proof-of-concept, the technique was applied to the analysis of mitochondria from hepatocytes exposed to silver nanoparticles. It was thus possible to identify mitochondria with higher concentration of Ag(I) ions compared to the surrounding cytosol. The versatility of the method makes it suitable to answer a large panel of biological questions, for instance related to metal homeostasis in biological organisms.

## Introduction

Metal ions are essential for Life. They are required for various processes such as redox-dependent enzymatic reactions, respiration or protein domain structuring. Besides, redox-active free metal ions can induce oxidative stress in cells. Therefore, living organisms tightly control metal ion uptake, intracellular trafficking, storage, and excretion. Although these complex mechanisms, termed homeostasis, are globally understood for Zn, Fe and Cu in Human, some molecular details remain unclear. For instance, FeS clusters biogenesis that occurs within the mitochondria is not completely understood (Beilschmidt et al., 2017). Moreover, disease-related metal overload or environmental exposure to metals or metal nanoparticles disrupt these mechanisms, altering cellular physiology. It is thus of paramount importance to be able to visualize trace element subcellular distribution, *i.e.* down to the organelle level for eukaryote cells.

Various approaches are used to study metal distribution within biological samples, with a large range of sensitivities and spatial resolutions (for review (Decelle et al., 2020)). For large field of view analysis with a moderate sensitivity, two methods exist: Laser-Induced Breakdown Spectroscopy (LIBS) and Laser Ablation Inductively Coupled Plasma Mass Spectrometry LA-ICP-MS (Busser et al., 2018; Greenhalgh et al., 2020; Wiemann et al., 2017). Their micrometer resolution makes them useful for the analysis of tissues. Nanometer resolution can be reached with secondary ion mass spectroscopy (SIMS) and Energy-dispersive X-ray spectroscopy (EDX) in an electron microscope (for a general review on the different methodologies (Dressler et al., 2018)). However, their sensitivity does not allow the detection of molecular species at a biologically relevant concentration. The method of choice to analyze the distribution of any element of interest with subcellular resolution, and sensitive enough to detect trace elements even in their soluble form, is X-ray fluorescence (XRF) imaging performed in a synchrotron X-ray nanoprobe (Fus et al., 2019; Hasna et al., 2019; Tardillo Suárez et al., 2020). Applied to nanotoxicology research, this technique provides the unique capability to detect both metallic nanoparticles and soluble metal ions in a biological matrix, as well as the native elements of the cell or tissue (Brown et al., 2018; Veronesi et al., 2019, 2016). As of today, high-resolution 2D elemental images have been most often obtained on entire cells, where the XRF signal from the whole thickness of the cell is summed up, losing the in-depth resolution and consequently the precise distribution within the cell down to the organelle level. In order to overcome this limitation, 3D tomographic XRF imaging should be used (Fus et al., 2019; Yuan et al., 2013). However, the long acquisition time needed to collect a number of projections that allow the accurate reconstruction of the cell volume prevents a routine use of this technique in biology. Possible solutions applied so far consist in either performing 2D XRF imaging over a large sample set, then confirming the findings on a few specimens by acquiring 3D data (Fus et al. 2019), or in coupling 2D XRF on whole cells with other techniques that allow the localization of subcellular compartments. Among the latter, it is worth mentioning organelle labelling followed by fluorescence microscopy (Carmona et al., 2019; Yuan et al., 2013), or TEM observations of cell sections (Veronesi et al., 2016). In this context, the possibility to perform correlative imaging of cell sections by EM and XRF is highly desirable. This requires the investigation of sample preparation and measurement strategies that allow the observation of the same specimen with the two techniques, in spite of the usually different experimental requirements applied to the two techniques individually. For instance, EM observations rely on heavy metal staining of the biological samples, whereas staining is not recommended for XRF imaging, in particular with metals that would interfere with the XRF signal from the native elements. A recent publication described a sample preparation strategy using only tannic acid for staining in order to avoid the addition of heavy metals and ensure the quality of XRF data (Kashiv et al., 2016). In spite of a more reliable detection of the distribution of native elements with respect to heavy metal staining, tannic acid staining results in a loss of contrast and in a limited ultrastructural resolution. Recently, we have been able to observe cell sections in both TEM and XRF imaging, in order to follow the entry of Ag(I) ions released from AgNPs into hepatocyte nuclei (Tardillo Suárez et al., 2020). Ag is a non-physiological metal, but the increasing use of AgNPs as biocides in consumer products and medical devices causes humans to be exposed and to accumulate Ag species in the organism. The binding properties of Ag(I) are similar to those of Cu(I), thus Ag(I) can replace Cu(I) in Cu-dependent enzymes, inhibiting them (Ciriolo et al., 1994). We also showed that Ag(I) can replace Zn(II) in the Zn-finger domain of specific transcription factors and thereof impair their activity (Kluska et al., 2020; Tardillo Suárez et al., 2020). Elemental XRF imaging was crucial to show the translocation of Ag(I) to the nucleus (Tardillo Suárez et al., 2020), suggesting that the observed impairment of nuclear functions originates from the binding of Ag(I) to transcription factors *in cellulo*. However, being the largest organelle in the cell, the nucleus was easy to identify by XRF imaging alone. In order to enhance the versatility of the method and be able to observe also smaller organelles, we need to obtain both the nanometer-resolution image of the interior of a cell and the corresponding elemental maps. This goal can be achieved by making use of correlative TEM and XRF microscopy, as described herein. As a proof of the interest of this methodology, we measured Ag(I) accumulation in the mitochondria of hepatocytes exposed to silver nanoparticles. We revealed that the Ag(I) concentration is higher in mitochondria compared to the surrounding cytosol, and we could observe a mitophagy process in case of Ag(I) overload. This case study illustrates the valuable contribution that the correlative imaging approach can bring to nanotoxicology research, by providing a tool to localize and quantify nanoparticles and trace elements at the organelle level. As such, this methodology can also be used to address questions related to metal homeostasis.

## Material and methods

### Cell culture

HepG2 cells were grown on labtek (Nunc) in MEM media supplemented with 10% Fetal Bovine Serum and 1% antibiotics-antimycotic at 37°C and in the presence of 5% CO_2_. Cells were exposed to 20 nm diameter citrate-coated AgNPs, cit-AgNP (Nanocomposix), or to 90 nm diameter PVP-coated AgNPs, PVP-AgNP (Sigma-Aldrich), at the indicated concentrations of total Ag and for the indicated times. AgNPs were previously characterized (Veronesi et al., 2016). Media and AgNPs were renewed every 24 hours.

### Sample preparation for correlative XRF and TEM imaging

Samples were prepared similarly to our recent work (Tardillo Suárez et al., 2020). At the desired timepoint, monolayers of HepG2 cells were fixed overnight at room temperature in a 1:1 ratio mixture of 4% paraformaldehyde, 0.4% glutaraldehyde in 0.2 M PHEM pH 7.2 and culture medium, washed in 0.1 M PHEM pH 7.2, and fixed for 30 minutes in 2% paraformaldehyde, 0.2% glutaraldehyde in 0.1 M PHEM pH 7.2, washed in 0.1 M PHEM pH 7.2 and post-fixed in 1% OsO_4_ in 0.1 M PHEM buffer for 1 h at room temperature. Cells were then dehydrated in graded ethanol series, and flat-embedded using an Epoxy Embedding Medium kit (Sigma-Aldrich). Sections (200 nm) were cut on a Leica UC7 ultra-microtome using a DiATOME 35° diamond knife and collected on 50 nm-thick Si_3_N_4_ grids (Oxford Instruments).

### Transmission electron microscopy

Digital images were obtained using a Tecnai G2 Spirit BioTwin microscope (FEI) operating at 120 kV with an Orius SC1000 CCD camera (Gatan).

### Nano-XRF data acquisition and analysis

XRF experiments were carried out at the state-of-the-art hard X-ray nanoprobe beamline ID16B at the ESRF (Martínez-Criado et al., 2016). The incoming photon energy was set at 29.6 keV and the beam was focused down to 55×65 nm^2^ using Kirkpatrick-Baez mirrors. The fluorescence emission from the sample was recorded using two 3-element Silicon Drift Detectors (SDD) arrays positioned at 13° from the sample. The photon flux on the sample was ~10^11^ photons/s. High resolution maps were recorded at room temperature, raster scanning the sample in the X-ray focal plane with 50×50 nm^2^ step size and 500 ms to 1 s dwell time/pixel.

Hyperspectral images were analyzed using the PyMCA software package (http://pymca.sourceforge.net/)(Solé et al., 2007). The detector response was calibrated over a thin film multilayer sample from AXO (RF8-200-S2453). XRF data were energy calibrated, normalized by the incoming photon flux, and batch-fitted in order to extract spatially resolved elemental concentrations, assuming a biological matrix of light elements and density of 1 g.cm^−3^ according to NIST standards (https://physics.nist.gov/cgi-bin/Star/compos.pl?matno=261).

## Results and discussion

### Implementation of correlative TEM and XRF microscopy

Recently, we successfully used XRF on cell sections for the analysis of Ag transformations and distribution within hepatocytes exposed to AgNPs (Tardillo Suárez et al., 2020). In the current study, the methodology was extended to be able to nail down the trace element content in the organelles of interest. Cells were grown on standard labtek supports that were used for chemical fixation using paraformaldehyde/glutaraldehyde. This is a good compromise that enables to preserve the intracellular content in comparison to alcohol-based fixation (Jin et al., 2017; Sanchez-Cano et al., 2017). Sample processing was based on classical cell resin-embedding preparation. However, to avoid parasite signals, all heavy metal used to stain and enhance the contrast in the sample (for instance U and Pb) have been removed, at the exception of OsO_4_. The latter is necessary to preserve the cellular ultrastructure by reacting with the lipids in the membranes. Therefore, OsO_4_ also provides contrast at the cellular membranes both in TEM and in XRF, thanks to the electron scattering and an X-ray absorption properties of the heavy Os atoms. The influence of the presence of OsO_4_ on the elemental content of fixed cells was estimated empirically, by comparing the elemental distributions of metals by means of XRF imaging in cells exposed to citrate-coated AgNPs for 72 h, then chemically fixed with or without OsO_4_ post-fixation. The average elemental composition per cell, estimated over three cells per condition, are reported in **Table S1**. The trace element contents in the cell area were measured, and showed that only in the sample post-fixed with OsO_4_ it is possible to distinguish basal Fe from the background, and that the native Cu is double with respect to the Os-free sample. It is not possible, instead, to determine the amount of native Zn in the presence of Os, because of the overlap of the X-ray emission lines of the two elements (**Fig. S1** and **Table S1**). Moreover, the cytosol concentration of molecular Ag(I) determined by XRF mapping is twice higher in the case of hepatocytes prepared with Os in comparison to cells prepared without, 560 +/− 90 ppm *versus 280* +/− 50 ppm, respectively (**Fig. S1**). Os fixation is therefore crucial to maintain the soluble complexes of biomolecules with metal ions inside cells and it was kept in our optimized sample preparation protocol. However, whenever the determination of Zn content is crucial for the scientific case under investigation, and if the X-ray beam energy is above the Os L_III_-edge excitation threshold (10.87 keV), another preparation protocol should be applied, by substituting OsO_4_ with another oxidant acting on lipids for example. Alternatively, the X-ray energy could be set between 9.66 keV (Zn K-absorption edge) and 10.87 keV, in such a way that only Zn atoms are excited, whereas Os atoms are not.

Thin sections produced by ultramicrotomy are deposited on Si_3_N_4_ EM grids comprising a square window of 500 μm wide and 50 nm thickness that are suitable for both EM and XRF imaging. The surface area is limited, which limits the number of cells in the window but facilitates the retrieval of the areas of interest. The Si_3_N_4_ EM grid is first scrutinized by TEM to identify cells with organelles and/or markers of interest inside. The position of the chosen cells is roughly found by the optical microscope on the ID16B beamline. Fast XRF mapping at low dose deposition is performed at low spatial resolution with 1×1 μm^2^ step size and 100 ms dwell time to confirm the location of the cells of interest by observing the Os signal. The latter can also provide shadows of the organelle and thus gives confidence in the retrieval of the area of interest. Fast scans also limit the radiation damage. In a second step, XRF mapping at high-resolution (down to 50×50 nm^2^ step size and with a dwell time of 500 ms per point or more) is performed to obtain high quality XRF data in terms of both spatial resolution and sensitivity. These maps are acquired on an area of the order of 10×10 μm^2^ in order to limit the acquisition time to a few hours.

A section of an HepG2 cell exposed for 6 hours to 50 μM cit-AgNPs was sequentially imaged by TEM (**Fig. 1A**) and then XRF (**Fig. 1B-C**). The low-resolution TEM image of the complete cell section revealed the different cellular compartments, for instance the largest that is the nucleus, which can also be identified in the XRF map of Os Lα since the nucleus is not penetrated by this metal species. As a proof of principle, this set of images enabled to observe electron dense objects by TEM (red and orange arrows) that can be attributed to Os deposits (orange arrows) and AgNPs in endosomes or lysosomes (red arrows) thanks to the extraction of the specific elemental maps of Os and Ag, respectively. Therefore, such a correlative use of TEM and XRF enables the simultaneous determination of the ultrastructure and of the elemental content of the interior of a cell. Such information can be retrieved also through TEM coupled to energy dispersive X-ray spectroscopy. However, the sub-ppm sensitivity of XRF is necessary to detect trace metal amounts of a few attograms (10-18 g) per pixel (Veronesi et al., 2016).

**Fig. 1.**
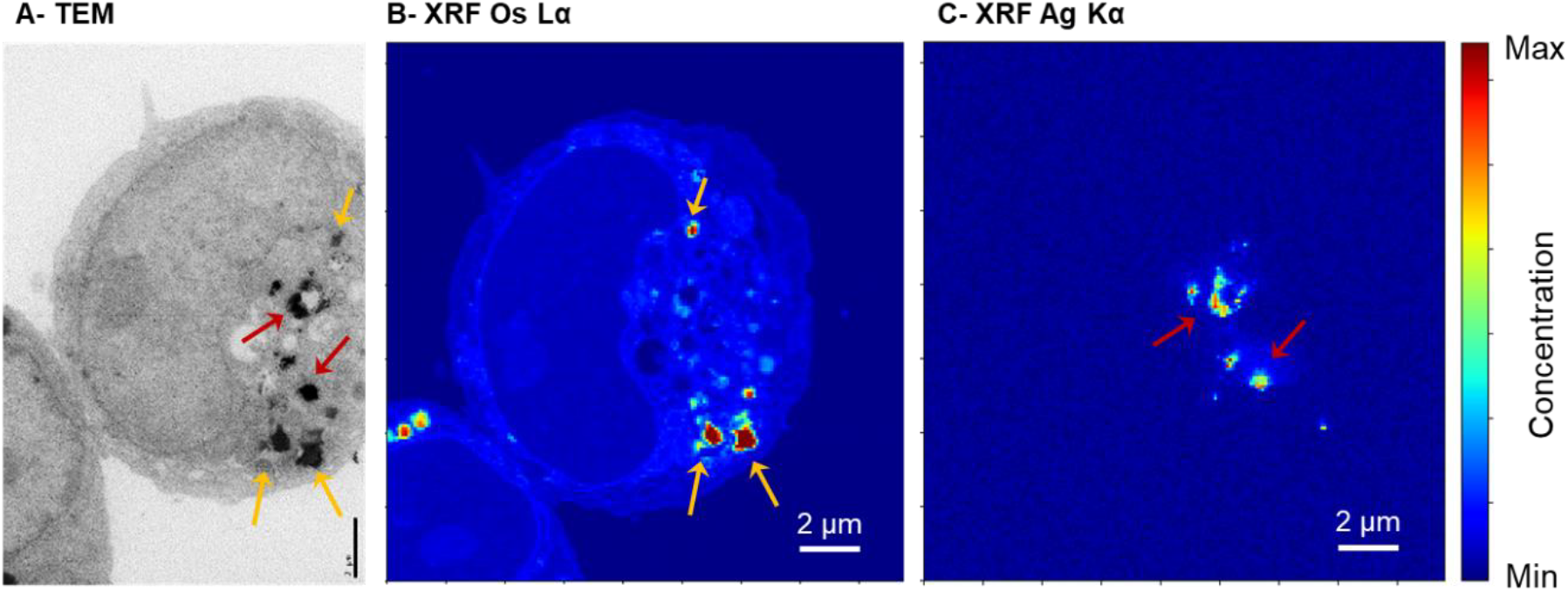
Correlative TEM and XRF for the identification of Os deposits *versus* AgNPs. Analysis of a 200 nm section of a HepG2 cell exposed to 50 μM cit-AgNPs for 6 h. (A) TEM image with orange and red arrows pinpointing areas of interest showed in XRF images. (B) Os and (C) Ag areal density maps in false colors for the same section. XRF acquisition was done with a pixel size of 100×100 nm^2^. All scale bars correspond to 2 μm.

### Sensitivity and spatial resolution of TEM-XRF in comparison to STEM-EDX

In a recent study, we considered two different chemical forms of silver: i) AgNPs, always found in endosomes and lysosomes, in their pristine form or partially transformed, and ii) molecular Ag(I)-thiolate species that were found, at least, in the cytosol or in nuclei (Tardillo Suárez et al., 2020). However, we expect to be able to disclose the distribution of cytosolic Ag between the different organelles such as mitochondria, endoplasmic reticulum or Golgi apparatus. It is frequently assumed that Ag(I) can accumulate into mitochondria by using the Cu(I) uptake pathway that is still undefined at the molecular level (Cobine et al., 2006). A section of HepG2 cells exposed to 25 μM PVP-AgNPs for 24 hours was inspected by TEM (**Fig. 2A**). We could evidence the nucleus surrounded by mitochondria (green rectangle in **Fig. 2A** and zoom-in in **Fig. 2B**). The area rich in mitochondria (red rectangle) was selected for further analysis by XRF, with the aim to compare Ag concentration in mitochondria and cytosol. These acquisitions were done at a resolution of 50×50 nm^2^ to obtain detailed images of mitochondria. Osmium enabled to confirm the location of mitochondria with respect to the nuclear membrane (**Fig. 2C**). Very interestingly, the Ag areal density map in false color (**Fig. 2D**) revealed a slightly higher concentration of Ag within mitochondria compared to the surrounding cytosol: 280 ppm in average inside mitochondria *versus* 160 ppm in the cytosol, after background subtraction and fitting of the XRF signal integrated over the two different cellular compartments. This proves that our analytical protocol allows for the subcellular mapping of trace elements.

**Fig. 2.**
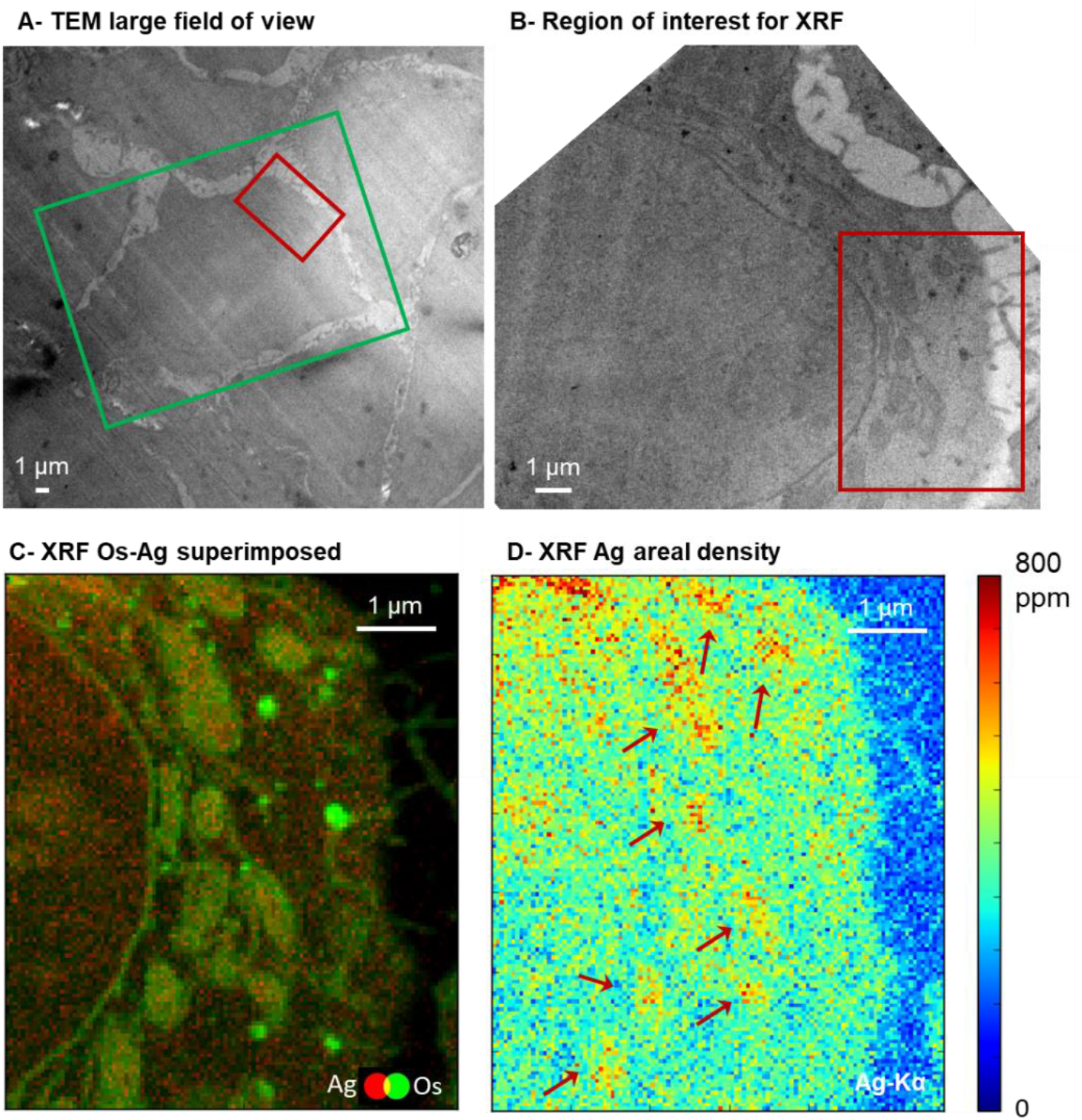
Ag(I) accumulation into mitochondria in HepG2 cells exposed to AgNPs. (A) Large TEM field of view with several cells visualized from a section of HepG2 cells exposed to 25 μM PVP-AgNPs for 24 hours. (B) Close-up view to the cell of interest, green rectangle in A. (C) Double color XRF map of the region of interest, red rectangle in A and B, with Os (green) and Ag (red) superimposed. (D) Ag areal density map represented in false-colors of the same region as in C. Red arrows pinpoint mitochondria. XRF acquisitions were done with a pixel size of 50×50 nm^2^.

Control cells, not exposed to AgNPs were also imaged using this correlative approach (**Fig. 3**). A field rich in mitochondria was identified by TEM (**Fig. 3A**, red rectangle) and then scanned by XRF (**Fig. 3B-C**). The Os map again revealed mitochondria shadows (**Fig. 3B**, red arrows) and the Ag map showed only noise signal (**Fig. 3C**). Altogether, our data showed that the correlative methodology developed can efficiently and simultaneously detect AgNPs, soluble silver species, and subcellular compartments. This enabled to detect minor differences in terms of trace element accumulation within a specific organelle, in this case the mitochondria.

**Fig. 3.**
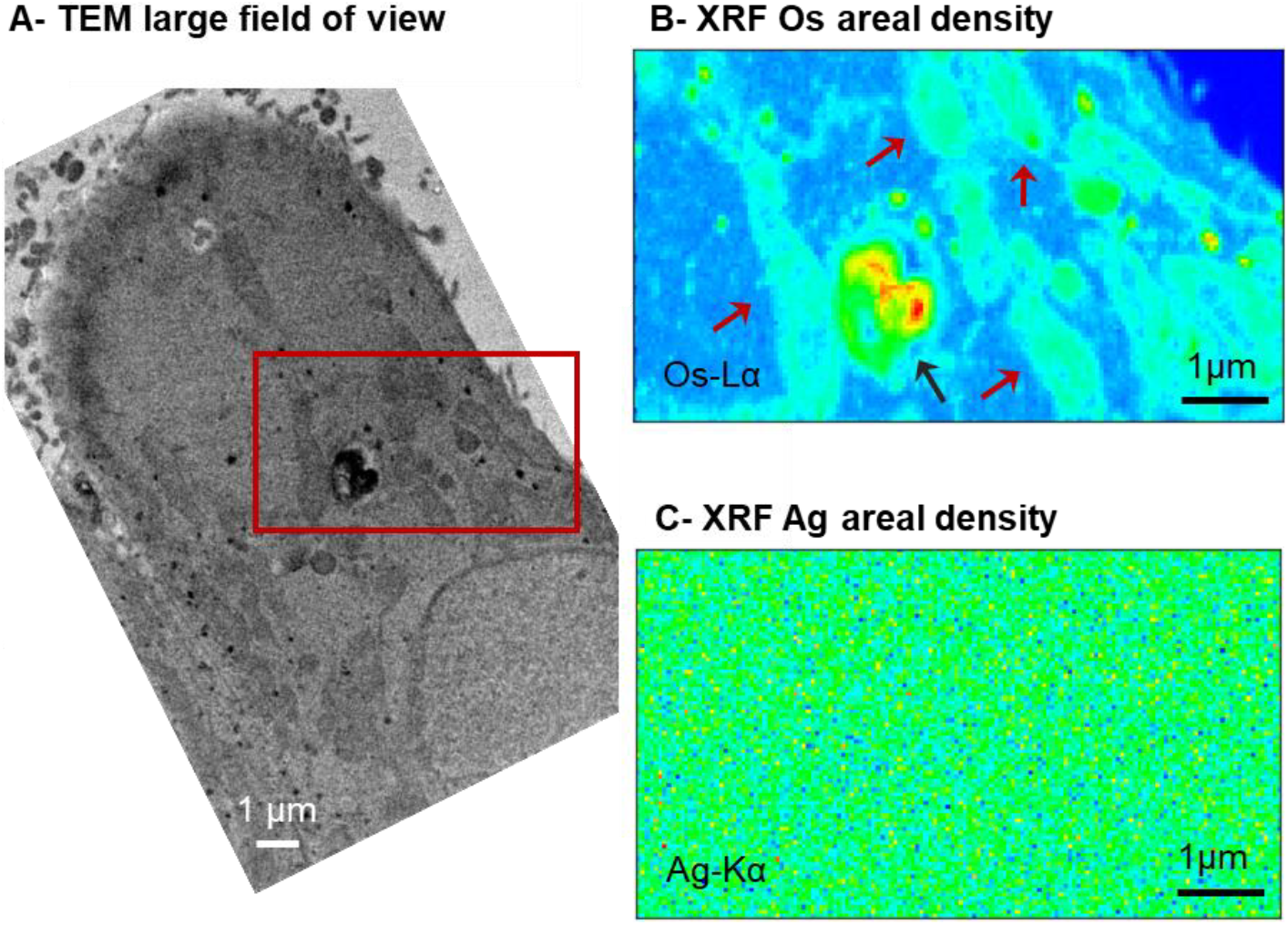
Control cells analysis. (A) TEM image of the region of interest in a section of HepG2 cells non-exposed to AgNPs. (B) Os and (C) Ag areal density maps in false colors for the same region of interest, red rectangle in A. Mitochondrial shadows are pinpointed with red arrows in B. Black arrow pinpoint a vesicle containing Os deposit as seen by TEM (A). XRF acquisition was done with a pixel size of 50×50 nm^2^.

To compare this multimodal approach with conventional STEM-EDX, a field comprising part of a nuclei, cytosol, a mitochondria (red arrow), and a lysosome containing AgNPs and transformed AgNPs (orange arrow) was imaged by STEM (**Fig. S2A-B**). EDX analysis showed that only electron dense spots within the lysosome present significant Ag signal (**Fig. S2C-D**), confirming that they were AgNPs or dissolving AgNPs. The mitochondria, the cytosol and the nucleus did not display Ag signal by EDX, while synchrotron nanoprobe XRF enabled not only to detect molecular Ag(I) species found in these three compartments but also to precisely quantify local Ag(I) concentrations and thus rank them according to their Ag(I) accumulation : the highest is in mitochondria, followed by the cytosol and finally the nucleus. Therefore, the combined ultrastructural imaging and trace element quantification obtained by the correlative TEM and XRF imaging approach provides additional information with respect to the STEM-EDX method.

### XRF offers a high dynamic range

Together with the sensitivity to low concentrations, the dynamical range is an important parameter in elemental imaging, in particular for elements that are present in both their crystalline form (*e.g.* nanoparticles) and as soluble molecular complexes. An example in biology is Fe, which is stored in ferritin in the mineral form and exists also as an enzyme co-factor. The dynamical range is also crucial in the determination of the intracellular fate of AgNP, where both crystalline and soluble silver need to be detected. Indeed, AgNPs are endocytosed and dissolved within endo-lysosomes into Ag(I) species that get distributed throughout the cell (Veronesi et al., 2016). XRF is quantitative over at least 5 orders of magnitude as seen in **Fig. 4**. XRF rendering using a linear scale ranging from 0 to 100000 ppm only highlights the presence of Ag-rich spots in vesicles, attributable to internalized AgNPs, together with a much weaker signal throughout the vesicles, most probably due to Ag(I) released from the NPs in the acidic lysosomal environment (**Fig. 4B**, **red arrows**). However, the visualization of Ag in the same area with a scale ranging between 0 and 1000 ppm highlights the presence of a much weaker signal (< 400 ppm) in the cell cytosol, not visible in the previous representations (**Fig. 4C**). Finally, the use of a logarithmic-scale representation of Ag distribution highlights the presence of all Ag species, regardless of the concentration, at a glance (**Fig. 4D**). Interestingly, the two last representations enables to visualize Ag molecular species that are at low concentrations. However, only the contrast of **Fig. 4C** reveals the capability of high resolution SR-XRF since in this representation both the spatial resolution and the sensitivity enable to confirm the different level of Ag molecular species in different organelles. Indeed, one can clearly see a very low Ag level for the nucleus, intermediate for the cytosol and the nucleolus, and two undefined vesicles show a significant accumulation of Ag (**Fig. 4C**, **white arrows**). This example demonstrates that XRF nano-imaging has the unique advantage to be able to detect simultaneously metallic nanoparticles and soluble metal complexes, providing high detection performances in a very large concentration range. It also highlights the importance of the choice of the visualization parameters to reveal the presence of all signals detected by the hyperspectral imaging technique. Finally, the high sensitivity and dynamic range of SR-XRF are interesting to correlate with EM ultrastructure imaging only because the spatial resolution progressed and became close enough to EM resolution.

**Fig. 4.**
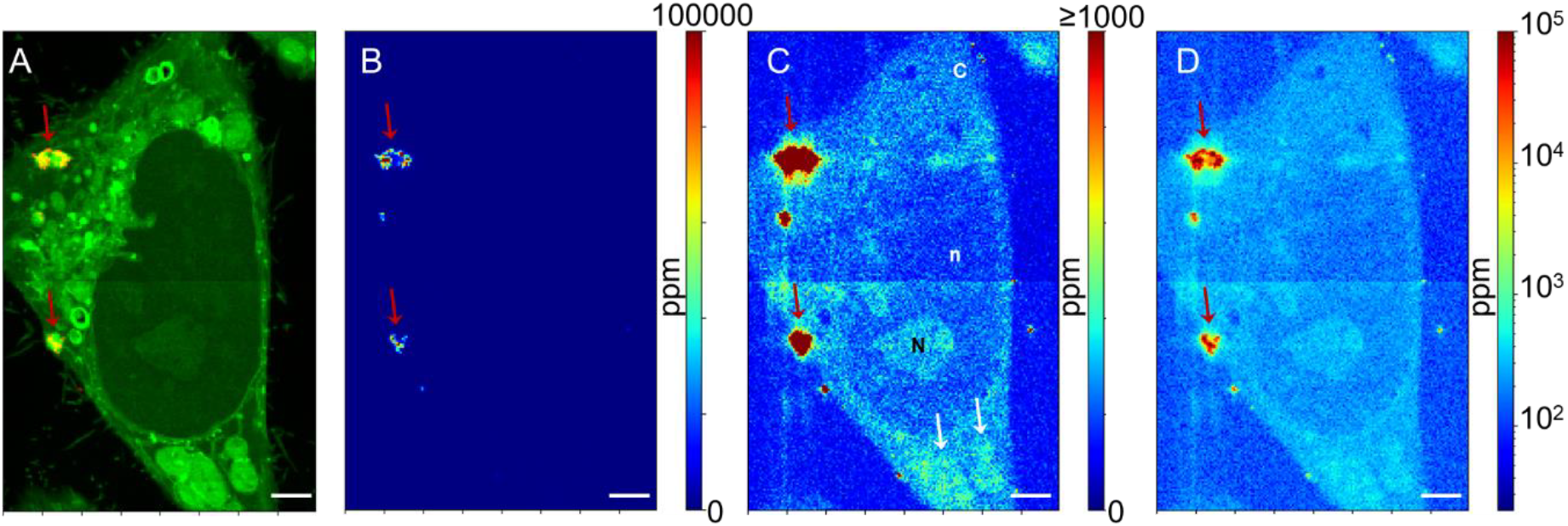
False-color representation of Ag distribution in hepatic cells exposed to 50 μM citrate-coated AgNPs for 72 h. (A) Os (green) and Ag (red) two-color map highlights the subcellular compartments and the presence of Ag-rich vesicles. (B) Ag distribution visualized using a linear color scale ranging between 0 and 100000 ppm. (C) Ag distribution visualized using a linear color scale ranging between 0 and 1000 ppm. (D) Red arrows pinpoint endo-lysosomes containing AgNP upon transformation and white arrows undefined vesicles with an accumulation of Ag molecular species. n, N and C stand for nucleus, nucleolus and cytosol, respectively. Logarithmic-scale representation of Ag distribution in the same area (0-100000ppm). Scale bars correspond to 2 μm.

### Catching events, the example of mitochondrial Ag overload-induced mitophagy

One main advantage of a single-cell, imaging-based analysis method compared to a bulk quantification approach is the possibility to identify specific events that can occur rarely, but that are interesting to understand completely a biological process. Similarly, an imaging method can also enable to dissect a whole process, by visualizing the cascade of events involved. In the data acquired on HepG2 cells exposed to 25 μM PVP-AgNPs for 72 hours, a mitochondrion located inside a double membrane vesicle was visualized by TEM (**Fig. 5A**, red arrow). This is clearly a mitochondrion that has been engulfed in an autophagosome, pinpointing the triggering of a mitophagy process (Ding and Yin, 2012) in this condition. The Ag areal density map of this region revealed a high Ag content in the mitochondria located in the autophagosome (**Fig. 5B**, red arrow). Indeed, the value was in the order of 5000 ppm, to be compared to the typical 200 to 700 ppm range observed in normal mitochondria in the different conditions tested. Therefore, the mitochondrion that underwent a mitophagy process accumulated 10-times more Ag than healthy mitochondria. This example highlights the importance of associating quantitative elemental distributions to high-resolution structural observations, as in the correlative TEM and XRF approach, since it revealed that the observed mitophagy is likely due to Ag overload in mitochondria. Current cellular data assumed similar Ag(I) molecular species uptake for a type of organelle and previous studies showed directly or indirectly a significant Ag(I) entry into mitochondria (Vest et al., 2013). However, our data showed a broad variability in silver amount accumulated in different mitochondria. Moreover, Ag hot spots were observed at the bottom of the XRF image in **Fig. 5B** (red pixels). This intensity is usually observed for AgNPs and/or transformed AgNPs, as suggested also by the TEM micrograph (**Fig. 5A**) that showed, in this area, electron dense spots typical of NPs. It is thus possible to imagine that mitochondria close to endosomes and/or lysosomes containing dissolving AgNPs are subjected to a massive accumulation of Ag that leads them to collapse and be degraded by mitophagy.

**Fig. 5.**
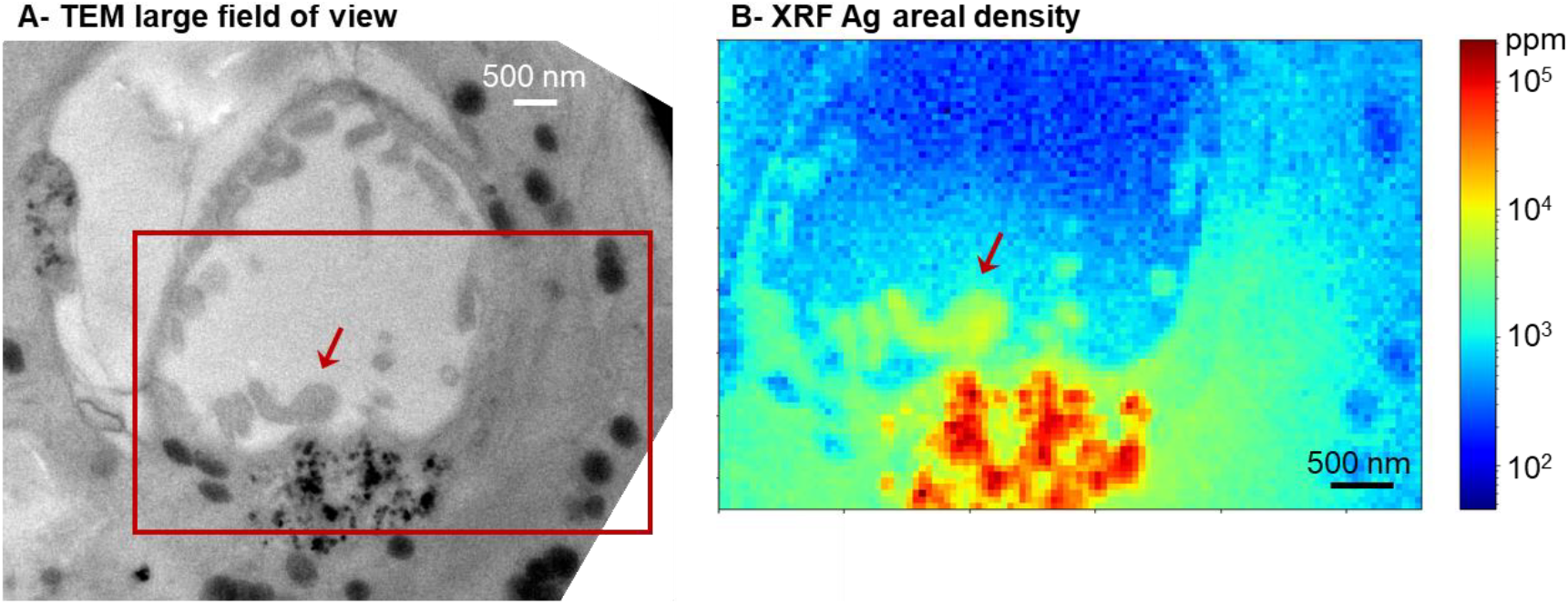
Ag-overload induce mitophagy. (A) TEM image from a section of HepG2 cells exposed to 25 μM PVP-AgNPs for 72 hours. (B) Ag areal density map in false colors of the region of interest, red rectangle in A. Red arrows pinpoint the same mitochondria. XRF acquisition was done with a pixel size of 50×50 nm^2^.*

## Conclusions

In the current study, we proposed the implementation of correlative TEM and hard X-ray nanoprobe XRF imaging performed on cell sections as prepared for classical cellular EM but with the only use of OsO_4_ as a heavy metal staining. Indeed, OsO_4_ is required both to protect trace elements from partial leakage out of the cell and to visualize cellular membranes where Os accumulates. This methodology has been optimized on 200 nm thickness samples. This thickness was chosen to provide, on the one hand, TEM images with enough details to identify the different organelles, and on the other hand, high enough XRF signal for trace element mapping. The combination of both enables to map elemental distributions at the organelle level. The quality of the data that can be obtained, as well as the sensitivity and the accuracy of the quantification, were evidenced in this study by the analysis of Ag(I) molecular species partitioning between mitochondria and the surrounding cytosol in hepatocytes that have been exposed to AgNPs. These experiments proved that, following AgNP dissolution into Ag(I), molecular species containing Ag(I) can enter mitochondria. The entry could be due to the recruitment of Ag(I) by the Cu(I) mitochondrial uptake pathway. Besides, we are inclined to think that the mitochondrial uptake of Ag(I) is an uncontrolled process, since a mitochondrion with ten times higher level than all others was found into an autophagosome. This event was observed close to a lysosome containing dissolving AgNPs. One can hypothesize that Ag(I) uptake is mainly driven by localization effect and no other control mechanism exists. Therefore, the Cu(I) trafficking pathway would not play a central role in Ag(I) organelle distribution and accumulation, as opposed to current models in the literature (Vest et al., 2013). All these data bring insight in the context of AgNP fate and toxicity in the liver, and demonstrate the usefulness of the method. The latter can be extended to the study of mechanisms involving other trace elements, such as iron or copper homeostasis, at the subcellular level. Herein, the chosen beam energy was 29.6 keV, which is optimal for Ag mapping but not for the detection of the lighter, native metals such as Cu or Zn, for which a lower excitation energy is required to obtain the highest sensitivity. Nonetheless, Os L edge can overlap with some elements of interest, for instance Zn. To overcome this limitation, the sample preparation could be optimized and OsO_4_ could be replaced by MnO_4_ or RuO_4_ for instance (Swartzendruber et al., 1995). In conclusion, the correlative TEM and XRF imaging approach presented here consists in optimizing the implemented use of the two techniques, commonly used separately, to obtain elemental distribution maps embedded into their ultrastructural context, which is crucial for the study of metal homeostasis mechanisms or to understand the fate of non-physiologic metals.

## Acknowledgements

The authors acknowledge the ESRF for providing beam time on the beamline ID16B-NA. This work used the platforms of the Grenoble Instruct-ERIC Center (ISBG: UMS 3518 CNRS-CEA-UGA-EMBL) with support from FRISBI (ANR-10-INSB-05-02) and GRAL (ANR-10-LABX-49-01) within the Grenoble Partnership for Structural Biology (PSB). The EM facility is headed by Guy Schoehn and supported by the Rhône-Alpes Region, the Fondation Recherche Medicale (FRM), the fonds FEDER and the GIS-Infrastrutures en Biologie Sante et Agronomie (IBISA). This research is part of the LabEx SERENADE (grant ANR-11-LABX-0064) and the LabEx ARCANE and CBH-EUR-GS (grant ANR-17-EURE-0003).

