## Supplement for "Correlative transmission electron microscopy and high-resolution hard X-ray fluorescence microscopy of cell sections to measure trace elements concentrations at the organelle level"

**Electronic Supplementary Information**

**Affiliations**

* Correspondence should be addressed to **Dr Giulia Veronesi**

or to **Dr Aurélien Deniaud**

| | [Fe] (ppm) | [Cu] (ppm) | [Zn] (ppm) | [Ag] (ppm) |
|---|---|---|---|---|
| No post-fixation | 14 ± 2 | 12 ± 5 | 15 ± 3 | 280 ± 50 |
| $OsO_4$ post-fixation | 125 ± 16 | 30 ± 20 | - | 560 ± 90 |

Table S1. Elemental compositions measured in HepG2 cells exposed for 72 h to 12.5 µM cit-AgNPs nanoparticles, then fixed as explained in the Methods section, by adding or not $OsO_4$ as a post-fixative.

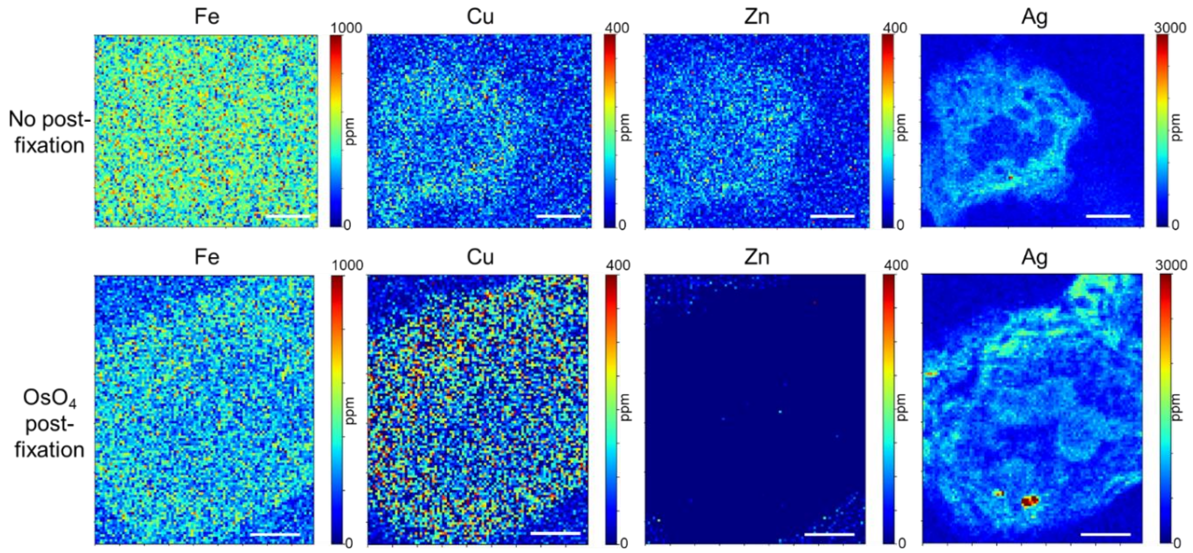


Fig. S1. Comparison of sample preparation with or without $OsO_4$. Fe, Cu, Zn and Ag concentration maps of 200 nm sections of HepG2 cells exposed to 12.5 µM cit-AgNPs for 72 h and prepared without $OsO_4$ (upper panels) and with $OsO_4$ (lower panels). Acquisitions were done with a pixel size of 100x100 $nm^2$. Scale bars correspond to 2 µm.

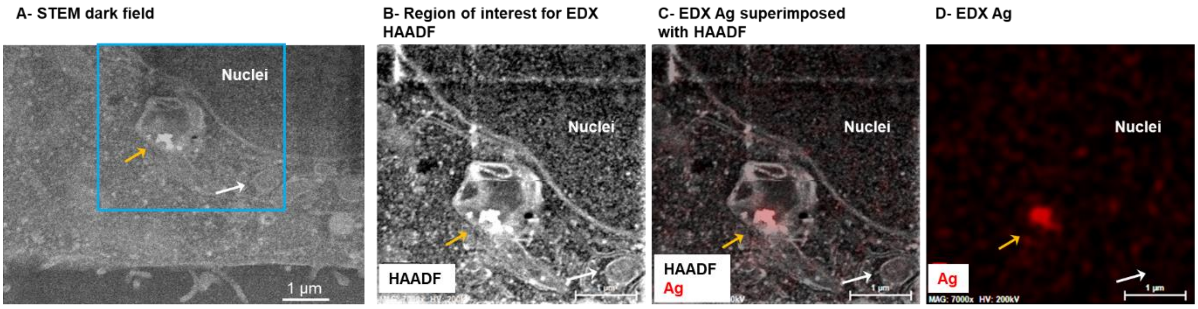


Fig. S2. AgNP fate analysed by STEM-EDX. (A) Large field of view STEM micrograph showing the presence of a lysosome containing NPs in contact with a nucleus in HepG2 cells exposed to AgNPs. (B) Dark-field STEM image of blue area in (A) at higher magnification. (C) Superimposition of dark-field image with Ag EDX map (red) and (D) EDX Ag map. Scale bars correspond to 1 µm.
